# Unveiling immune interference: How the dendritic cell response to co-infection with *Aspergillus fumigatus* is modulated by human cytomegalovirus and its virokine CMVIL-10

**DOI:** 10.1101/2025.05.15.654194

**Authors:** Linda Heilig, Lydia Bussemer, Lea Strobel, Kerstin Hünniger-Ast, Oliver Kurzai, Arnhild Grothey, Lars Dölken, Kerstin Laib Sampaio, Gianni Panagiotou, Alexander J. Westermann, Hermann Einsele, Sebastian Wurster, Sascha Schäuble, Jürgen Löffler

## Abstract

Human cytomegalovirus (HCMV) is a master of immune evasion and a potent modulator of the human immune system. The best-characterized mechanism employed by HCMV to suppress host immunity is the production of a viral interleukin-10 homolog (_CMV_IL-10). While _CMV_IL-10 is known to suppress immune responses and promote viral persistence, its capacity to promote increased susceptibility to co-infecting pathogens like *Aspergillus fumigatus* remains unknown. Therefore, we studied the impact of wild-type (WT) HCMV (strain TB40 BAC4), a _CMV_IL-10-deficient HCMV mutant (Δ*UL111A*), and recombinant _CMV_IL-10 on the immune activity of monocyte-derived dendritic cells (moDCs) during co-infection with *A. fumigatus*. Using a combination of transcriptomic and phenotypic readouts, our data revealed a strong and time-dependent immuno-paralytic effect of HCMV by suppressing pathogen recognition pathways, cytokine production, DC maturation, and expression of genes that are essential for host defense and tissue repair. Although infection with Δ*UL111A* lacking _CMV_IL-10 led to stronger expression of type I interferons, IFN-γ-inducible chemokines, and proinflammatory cytokines than WT infection, interference with antifungal immune defense and fungal clearance during co-infection was largely similar between both strains. The limited effect of _CMV_IL-10 on antifungal immune defense persisted even after prolonged pre-exposure of DCs to the recombinant virokine. In summary, although _CMV_IL-10 contributes to shaping an anti-inflammatory environment, HCMV’s suppression of antifungal immunity appears to be multifactorial, with _CMV_IL-10 alone playing a rather subtle role in altering DC responses to *A. fumigatus* during viral-fungal co-infection.

**Importance:** Human cytomegalovirus (HCMV) is a highly prevalent herpesvirus that establishes lifelong latency and frequently reactivates in immunocompromised individuals, including hematopoietic stem cell transplant recipients. Reactivation not only causes direct disease but also increases the risk of secondary infections, such as invasive pulmonary aspergillosis caused by *Aspergillus fumigatus*. Specifically, studies estimated that about 6-25% of critically ill HCMV-positive patients develop HCMV-associated pulmonary aspergillosis. However, the mechanisms by which HCMV creates a permissive environment for fungal superinfection remain poorly understood. HCMV encodes a viral homolog of interleukin-10 (_CMV_IL-10), which mimics host IL-10 and elicits potent immunomodulatory activity. Here, we show that _CMV_IL-10 dampens specific anti-viral responses, DC activation, and cytokine signaling. However, HCMV-mediated impairment of fungal control in co-infection settings occurred largely independent of _CMV_IL-10 expression. These findings suggest that HCMV undermines antifungal defenses through multifactorial mechanisms beyond _CMV_IL-10, highlighting the need for targeted strategies to restore immune function in high-risk patients.

## Introduction

Human cytomegalovirus (HCMV) establishes lifelong latency, with reactivation posing serious risks for immunocompromised patients, especially those undergoing solid organ or allogeneic stem cell transplantation (alloSCT) (1, 2). In alloSCT recipients, immune suppression and delayed immune recovery substantially increase the risk of reactivation (1, 3, 4). HCMV reactivation in these patients can progress to severe end-organ disease (EOD), affecting various organ systems including the lungs (2, 5) and contributing to significant mortality and morbidity (5).

Beyond the immediate risk of HCMV-induced EOD, reactivation can increase susceptibility to secondary pulmonary infections, particularly fungal pneumonias (1, 3, 6). Invasive pulmonary aspergillosis (IPA), primarily caused by *Aspergillus fumigatus*, is the most common fungal superinfection in patients with HCMV reactivation (3, 7). Concerningly, cohort studies estimated that about 6-25% of critically ill HCMV-positive patients develop HCMV-associated pulmonary aspergillosis (8–10). Co-occurring IPA in patients with HCMV disease is associated with increased mortality and poor prognosis (3, 6, 11). Notably, HCMV infection has been identified as an independent risk factor for the development of IPA in these patients (3, 12–14), suggesting inter-kingdom synergies between the two pathogens that create a permissive environment for fungal colonization and invasion (8, 15–17).

The complex interplay between viral and fungal co-pathogens relies on pleiotropic alterations of host defense, including epithelial barrier damage, impaired innate immune responses, and dysregulated cytokine signaling (15, 18–20). For instance, we previously found that HCMV suppresses antifungal immunity by counteracting *A. fumigatus*-induced activation of NF-κB (nuclear factor κB) and NFAT (nuclear factor of activated T cells) cascades, thereby attenuating the induction of proinflammatory cytokines such as IL-1B (16). In turn, *A. fumigatus* co-infection impairs viral clearance by disrupting antiviral signaling pathways like cGAS-STING signaling, leading to reduced production of IFNB, CXCL10, and CXCL11 (16).

Viral immune evasion mechanisms or virulence factors might play a role in shaping antifungal immunity (15, 21). HCMV employs sophisticated strategies to manipulate host immunity, evade detection, and sustain persistent infection (2, 22). The best-characterized mechanism employed by HCMV to suppress host immunity is the production of a viral interleukin-10 homolog (_CMV_IL-10) encoded by the *UL111A* gene (23–25). This virokine mimics the immunosuppressive effects of human cellular IL-10 and prevents NF-κB pathway activation, thereby suppressing proinflammatory cytokine production and antigen presentation (24, 26). Specifically, _CMV_IL-10 was shown to dampen activation, maturation, and function of dendritic cells (DCs), which are critical for orchestrating antifungal immunity (27–29). Levels of IL-10 are elevated during both the active and latent phases of HCMV infection (30–32). This mechanism likely plays a critical role in facilitating the establishment and long-term persistence of HCMV (30).

In addition to promoting latent viral infection (33, 34), _CMV_IL-10 may also increase susceptibility to co-pathogens like *A. fumigatus*. However, the immunomodulatory capacity of _CMV_IL-10 in immune cell-mediated cross-kingdom interactions with co-pathogens and the specific significance of _CMV_IL-10 in the *A. fumigatus*-DC relationship are largely unexplored. Therefore, we confronted human monocyte-derived dendritic cells (moDCs) with HCMV reference strain TB40 BAC4 (BAC4), an HCMV mutant lacking _CMV_IL-10 (Δ*UL111A*), or recombinant _CMV_IL-10, with and without subsequent *A. fumigatus* challenge (Figure 1). To assess moDC activation and modulation of the immune response upon single infection and co-infection, we employed a comprehensive approach including RNA sequencing (RNA-seq), flow cytometry, multiplex cytokine assays, and time-lapse microscopy (Figure 1). While confirming a severe immuno-paralytic effect of HCMV with profound suppression of genes and pathways that are crucial for moDC activation, anti-*Aspergillus* defense, and tissue repair, our findings suggest a rather subtle role of _CMV_IL-10 in this clinically important immune-mediated inter-kingdom synergy.

**Figure 1:**
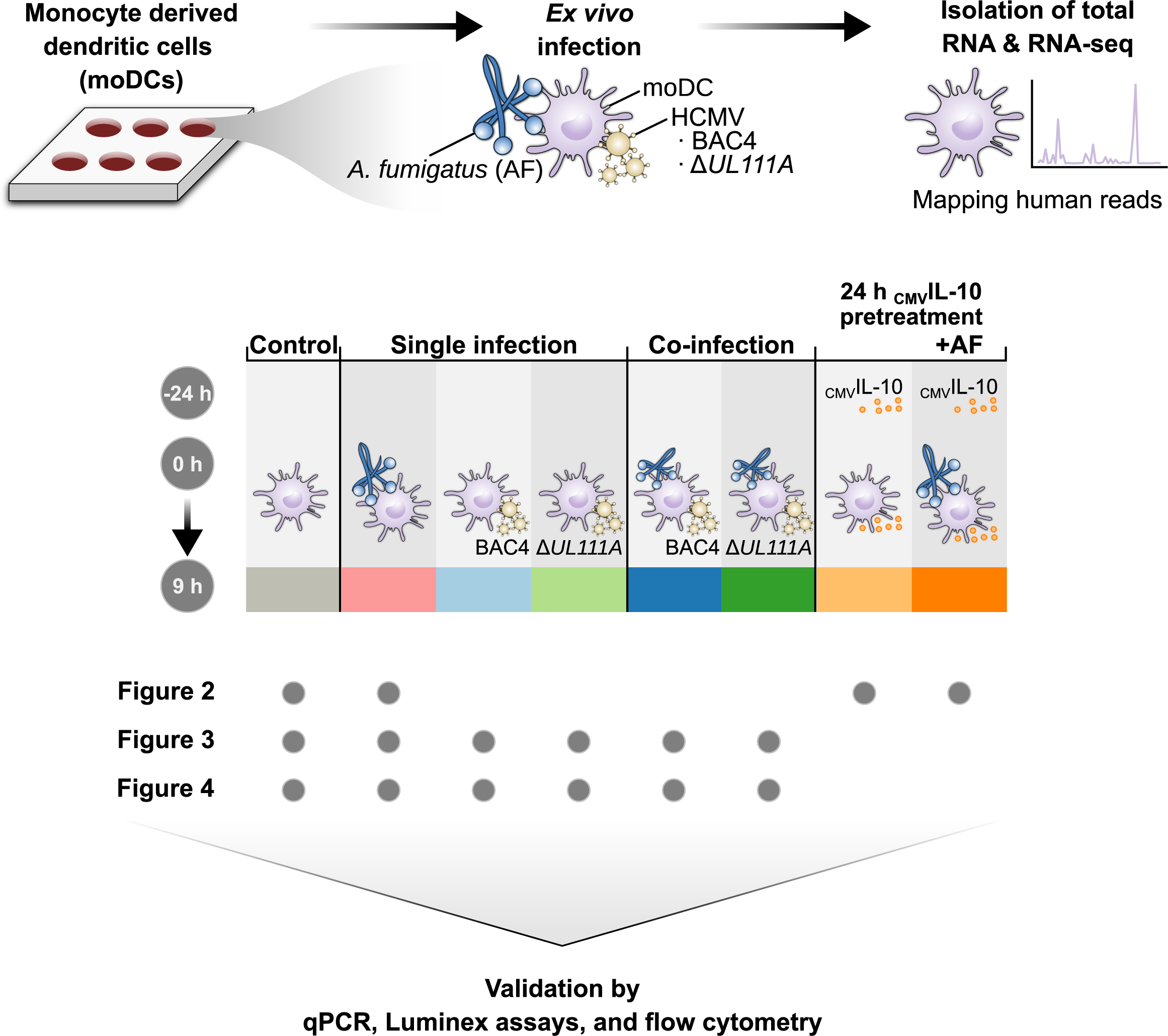
Schematic study design. Study design illustrating experimental setup of _CMV_IL-10 pretreatment and *A. fumigatus* (AF) infection. In addition, single (BAC4, Δ*UL111A*, AF) and co-infection (AF+ BAC4, AF + Δ*UL111A*) experimental setups are displayed.

## Results

### Recombinant _CMV_IL-10 alters expression of immune-related genes but does not specifically interfere with anti-*Aspergillus* responses of moDCs

At first, we sought to determine the impact of isolated _CMV_IL-10 on DC biology and responses to subsequent *A. fumigatus* challenge. As our previous study of the immune interference between HCMV and *A. fumigatus* had shown dynamic changes in gene expression within few hours post-infection (16), we focused on early changes in DC biology. Thus, moDCs were incubated with or without 50 ng/mL of recombinant _CMV_IL-10 for 24 h prior to challenge with *A. fumigatus* (27). Immuno-biological changes in moDCs were then studied by host-directed RNA-seq. Transcriptomes of uninfected cells were compared to transcriptional data from *A. fumigatus* single infection with or without _CMV_IL-10 pretreatment.

Principal component analysis (PCA) of transcriptional responses showed a clear separation of all four experimental conditions along the first two principal components (Figure 2A). Interestingly, combined _CMV_IL10 and *A. fumigatus* challenge formed a distinct cluster with a significant distance from both the control and single-treatment groups, suggesting additive or synergistic effects in gene regulation.

**Figure 2:**
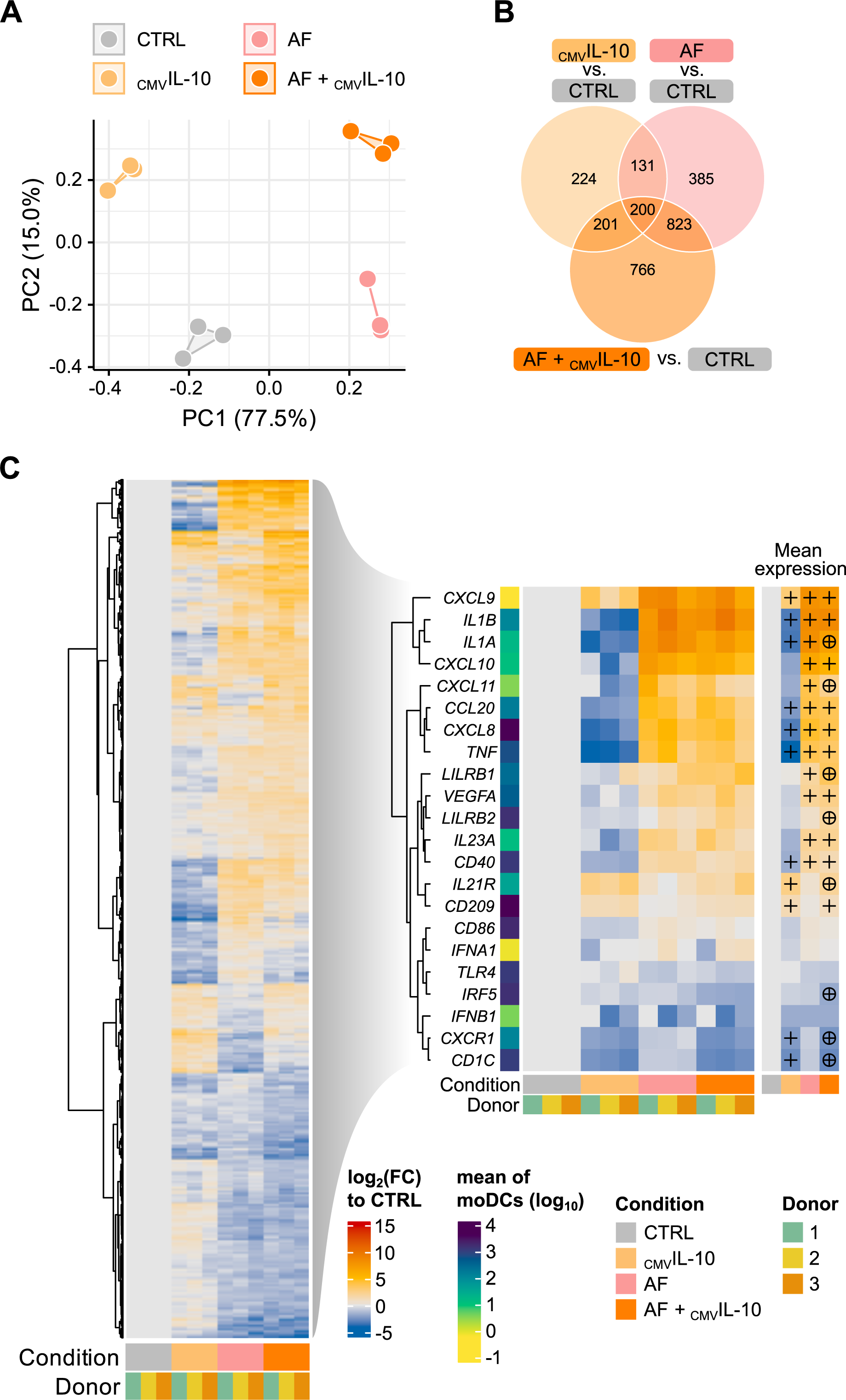
High concentrations of CMVIL10 attenuate several moDC effector responses to *A. fumigatus*. (A) Principal component analysis (PCA) comparing transcriptomes of unchallenged moDCs (CTRL) with expression profiles of moDCs exposed to _CMV_IL-10, *A. fumigatus* (AF), or both (AF + _CMV_IL-10). (B) Venn diagram summarizing shared and condition-specific differentially expressed genes (DEGs) across multiple comparisons. (C) Heatmap displaying DEGs in response to single infections with _CMV_IL-10 or AF, and co-stimulation. Left: DEGs associated with infection response. Right: Selected immune-related DEGs. Significant differential expression against “CTRL” (symbol: +) or “AF” (symbol: ⭘) according to DESeq2 analysis (adjusted p ≤ 0.05, log2(|FC|)≥1). N = 3 independent donors.

While single stimuli, i.e., either _CMV_IL-10 pre-incubation or *A. fumigatus* infection, led to unique sets of differentially expressed genes (224 and 385 genes, respectively), combination of the two challenges elicited a total of 766 unique DEGs (Figure 2B). A substantial number of DEGs (200 + 823 genes), however, were shared between *A. fumigatus* single infection and the combination of *A. fumigatus* together with _CMV_IL-10, also indicating a pronounced *A. fumigatus* specific impact irrespective of _CMV_IL-10 pretreatment.

To further dissect these transcriptomic differences, we focused on a comparison of immune-relevant transcripts across the experimental conditions yielding 2808 DEGs in total (Figure 2C). Notably, substantial differences in cytokine and chemokine gene expression were observed. Infection with *A. fumigatus* strongly induced VEGFA and proinflammatory markers such as *CXCL8*, *CXCL9*, *CXCL10*, *CXCL11*, *TNF*, *CCL20*, *IL1A,* and *IL1B* compared to control (Figure 2C, right part; Figure S1A und S1B). Inversely, _CMV_IL-10 pretreatment of moDCs without *A. fumigatus* infection resulted in a pronounced suppression of these genes. Interestingly, pretreatment with _CMV_IL-10 significantly attenuated *A. fumigatus*-induced upregulated transcription and/or secretion of some pro-inflammatory cytokines, such as IL-1A, TNF-α and CXCL11, while augmenting *A. fumigatus*-induced upregulation of *LILRB1* and *LILRB2* transcription (Figure 2C, right part; Figure S1A and S1B). Furthermore, _CMV_IL-10 slightly amplifies the *A. fumigatus*-induced secretion of cellular IL-10, further corroborating that _CMV_IL-10 exerts an immunosuppressive effect on moDCs (Figure S1A).

_CMV_IL-10 pretreatment also altered expression of moDC cell surface markers. For instance, _CMV_IL-10 induced significant upregulation of *CD209*, supporting HCMV entry (Figure 2C, right part; Figure S1C). In contrast, expression of *CD40* (transcriptional level) and CD86 (transcriptional and protein level), key molecules for antigen presentation, was reduced after _CMV_IL-10 pretreatment. These trends persisted when comparing combined _CMV_IL-10 and *A. fumigatus* challenge versus *A. fumigatus* single challenge (Figure 2C, right part; Figure S1B and S1C).

Overall, these findings support the immunosuppressive capacity of _CMV_IL-10 on various moDC effector functions. However, even after 24-hour pretreatment with high concentrations of recombinant _CMV_IL-10, its specific impact on *A. fumigatus*-induced responses remained relatively modest. Instead, our findings suggest largely additive effects of the two challenges rather than a specific interference of _CMV_IL-10 in the co-infection setting.

### DC responses to single and co-infections with HCMV and *A. fumigatus* cluster by infection type rather than viral _CMV_IL-10 expression

Next, we assessed the direct impact of HCMV Δ*UL111A*, lacking _CMV_IL-10 on the anti-*Aspergillus* response during co-infection. Consistent with prior data (16), PCA of moDC transcriptomes revealed distinct clusters based on infection type, with clear separation of cells infected solely with *A. fumigatus*, HCMV, or both pathogens (Figure 3A). In contrast, single or co-infections with BAC4 or Δ*UL111A* did not segregate by viral variant. Instead, BAC4 and Δ*UL111A* single infections clustered closely, as did their respective co-infection conditions (Figure 3A).

**Figure 3:**
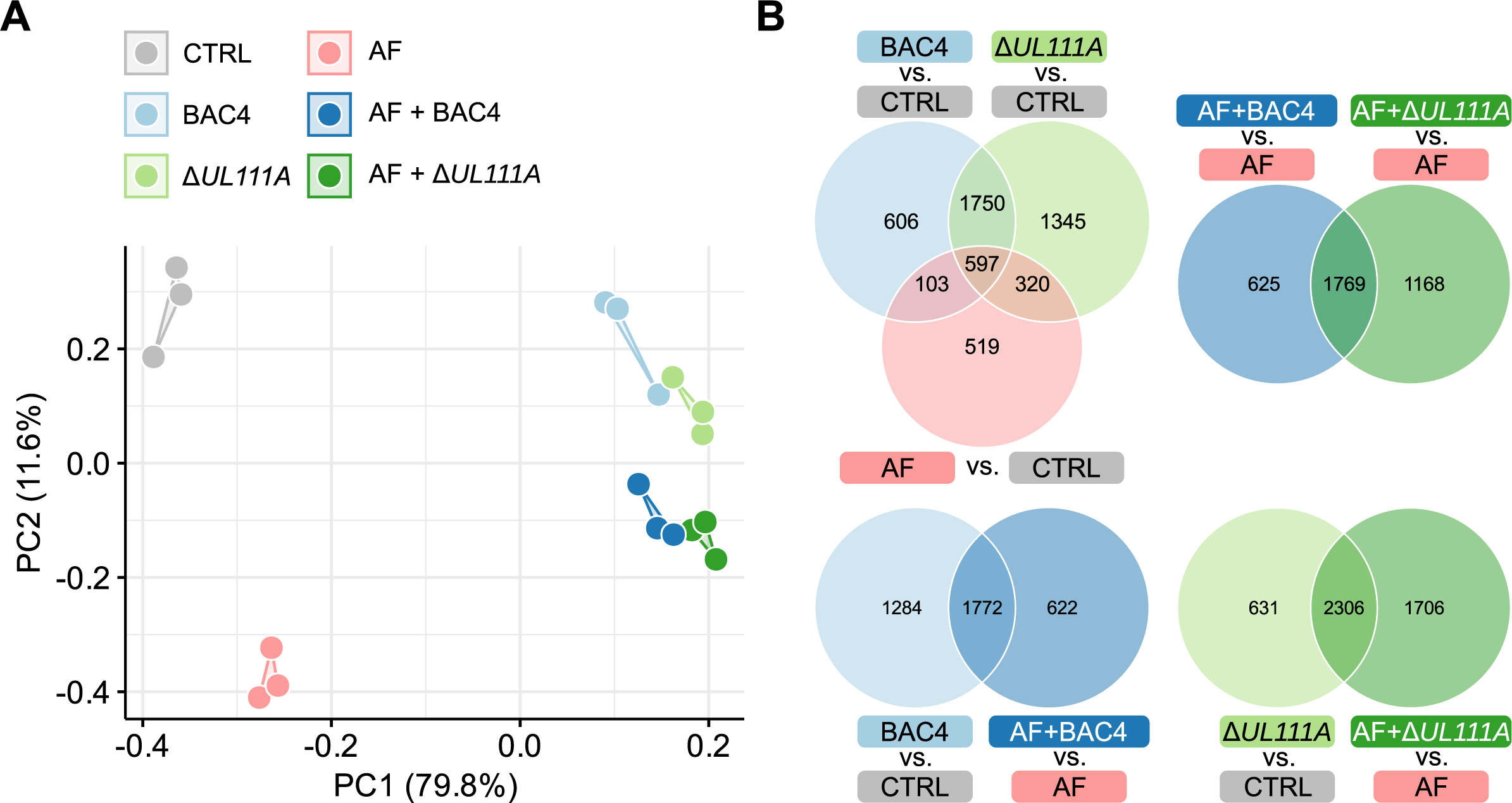
moDC expression profiles differ by infection type, with a rather subtle impact of _CMV_IL-10 expression. (A) Principal component analysis (PCA) comparing transcriptomes of moDCs (CTRL) with expression profiles of moDCs infected with *A. fumigatus* (AF), HCMV BAC4 or Δ*UL111A*, or both HCMV and AF (AF + BAC4, AF + Δ*UL111A*). (B) Venn diagrams summarizing shared and condition-specific differentially expressed genes (DEGs) across multiple comparisons. (A-B) N=3 independent donors.

Comparative analysis of the transcriptomes from single-infected moDCs (*A. fumigatus*, BAC4, Δ*UL111A*) versus uninfected controls identified 597 DEGs shared across conditions, along with genes uniquely expressed in each single-infection condition (BAC4 606 genes, Δ*UL111A* 1,345 genes, and *A. fumigatus* 519 genes) (Figure 3B). moDCs infected with HCMV BAC4 or Δ*UL111A* but not with *A. fumigatus* shared a high number of DEGs (1,750 genes) compared to uninfected moDCs. Likewise, direct comparison of the two co-infection conditions revealed a substantial number of 1,769 shared DEGs compared to *A. fumigatus* single-infection, regardless of viral _CMV_IL-10 expression. Collectively, these findings suggest a predominance of shared response patterns in co-infection settings with *A. fumigatus* and either WT or _CMV_IL-10-deficient HCMV.

### Viral _CMV_IL-10 subtly impacts HCMV-induced suppression of antifungal responses during co-infection

Next, we performed an analysis of DEGs associated with infection response (Figure 4A, left part, n = 6,435 total DEGs). Specifically, we focused our analysis on genes involved in immune and inflammatory processes, including genes for chemokines (*CXCL9, CXCL10, CXCL11, CXCL8*), pro-inflammatory cytokines (*IL1A, IL1B, TNF*), immune receptors (*CD40, CD86, IL21R, CXCR1*), and pattern recognition receptors (*TLR4, LILRB1, LILRB2*) (Figure 4A, right part).

**Figure 4:**
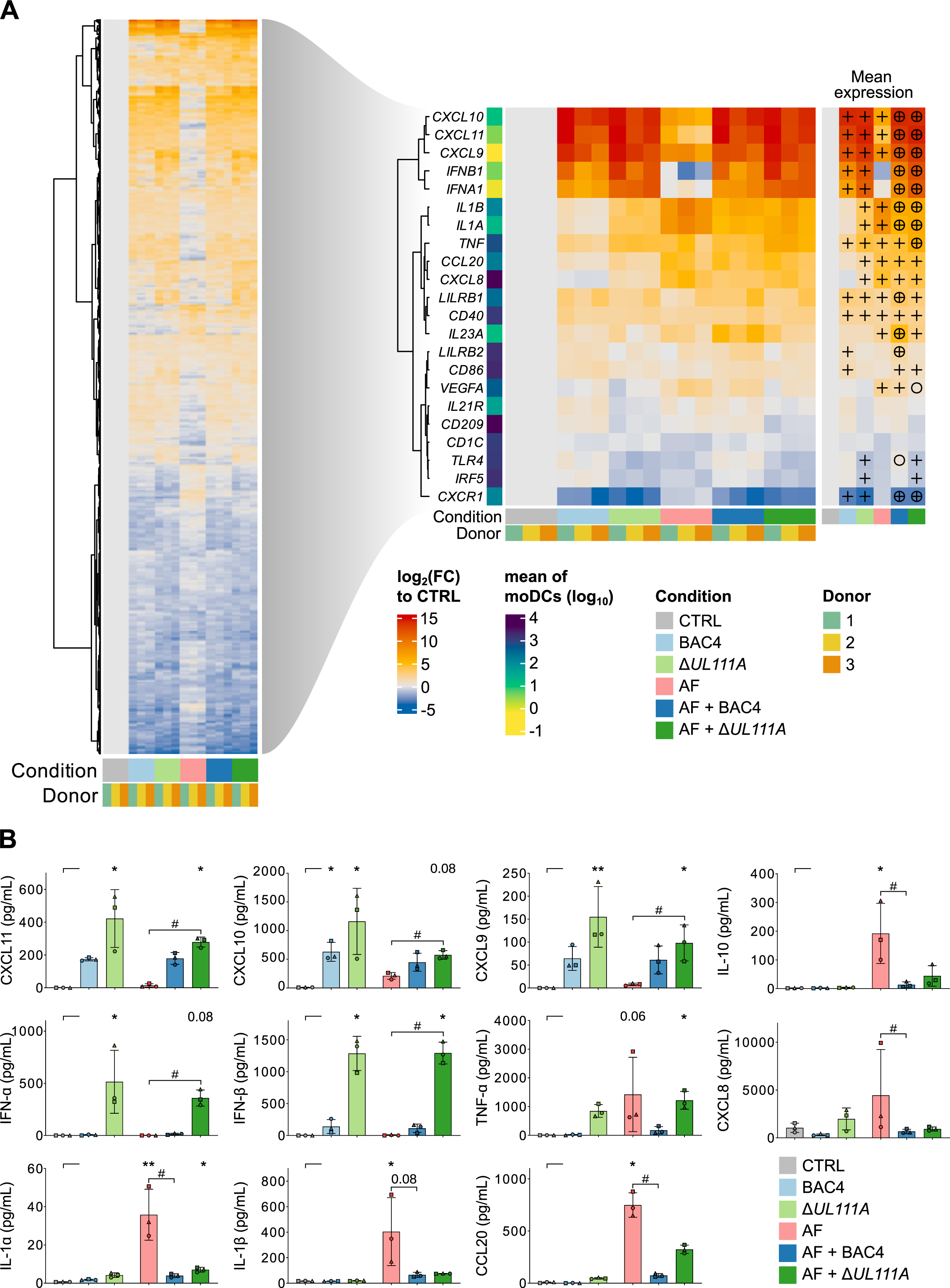
Despite differences in antiviral protein secretion, viral _CMV_IL-10 expression only subtly impacts HCMV-induced suppression of antifungal responses during co-infection. (A) Heatmap displaying differentially expressed genes (DEGs) in response to single infections with HCMV BAC4, Δ*UL111A*, or *A. fumigatus* (AF), as well as co-infection with both pathogens. Left: DEGs associated with infection response. Right: Selected immune-related DEGs across 6 tested conditions. Significance of pairwise comparison against “CTRL” (symbol: +) or “AF” (symbol: ⭘) is indicated. (B) Comparison of cytokine and chemokine release of uninfected moDCs (CTRL) and those challenged with single infection (BAC4, Δ*UL111A*, AF) or co-infection (AF+ BAC4, AF + Δ*UL111A*) for 9 h. (A-B) N = 3 independent donors. (A) Statistical significance according to DESeq2 analysis (adjusted p ≤ 0.05, log2(|FC|)≥1). (B) Columns and error bars indicate means and standard deviations, respectively. Friedman test with Dunn’s multiple comparisons test versus CTRL (asterisks). In addition, single *A. fumigatus* infection (AF) was compared to co-infection (AF + BAC4, AF + Δ*UL111A*) using Friedman test with Dunn’s multiple comparisons test versus AF (hash signs). */# p < 0.05, **/## p < 0.01.

We found significant upregulation of genes encoding IFN-γ-inducible chemokines (*CXCL9, CXCL10, CXCL11*) in both HCMV-infected conditions (Figure 4A, right part). These patterns were also reflected at the protein level (Figure 4B).

MoDCs exposed to HCMV strain Δ*UL111A* lacking _CMV_IL-10 displayed a trend toward stronger induction of CXCL9-11 than those infected with the WT strain (BAC4), especially on protein level (Figure 4A, right part; Figure 4B). Notably, the inhibitory receptors LILRB1 and LILRB2, which restrain antigen presentation and inflammatory signaling (35, 36), were upregulated in response to HCMV, with LILRB2 being more prominently induced following BAC4 infection. The expression of LILRB1 and LILRB2 was further amplified under co-infection conditions with BAC4 and *A. fumigatus* (Figure 4A, right part; Figure S2A). While infection of moDCs with either HCMV variant enhanced the expression of TLR3, cGAS, and hSTING (Figure S2B), the _CMV_IL-10 deficient strain elicited more robust transcription and secretion of type-I interferons (IFNA/B) (Figure 4A, right part; Figure 4B).

Consistent with the response to A. fumigatus in combination with _CMV_IL-10 (Figure 2C), pro-inflammatory genes such as *IL1A*, *IL1B*, *CCL20*, *VEGFA,* and *CXCL8* were predominantly upregulated following *A. fumigatus* infection, a trend that was confirmed at the protein level (Figure 4A, right part; Figure 4B; Figure S2A). Additionally, *A. fumigatus* induced expression of CXCL9, CXCL10, and CXCL11, especially on RNA level.

While co-infection with either HCMV strain markedly suppressed IL-1A and IL-1B transcription and release, few *A. fumigatus*-induced cytokines showed distinct modulation depending on the HCMV strain’s capacity to produce _CMV_IL-10. Specifically, *A. fumigatus*-induced expression and/or release of TNF-α, CCL20, and VEGFA were more potently attenuated by the _CMV_IL-10-producing WT strain (BAC4) than the Δ*UL111A* mutant (Figure 4A, right part; Figure 4B; Figure S2A).

This observation prompted us to explore transcription and release of cytokines that are important for DC activation and fungal clearance after prolonged pre-exposure of moDCs to HCMV (Figure 5). Pre-incubation with either HCMV variant for 24 h or 72 h before infection with *A. fumigatus* significantly and time-dependently suppressed gene expression and secretion of key cytokines such as CXCL8, VEGFA, IL-1α, IL-1β, IFN-γ, and TNF-α (Figure 5). However, none of these effects was influenced by the HCMV variant’s _CMV_IL-10 status. Collectively, these findings are consistent with a strong immuno-paralyzing effect of HCMV effect through downregulation of key antifungal responses while pointing to a limited role of _CMV_IL-10 expression in modulating the DCs’ antifungal response in a co-infection setting.

**Figure 5:**
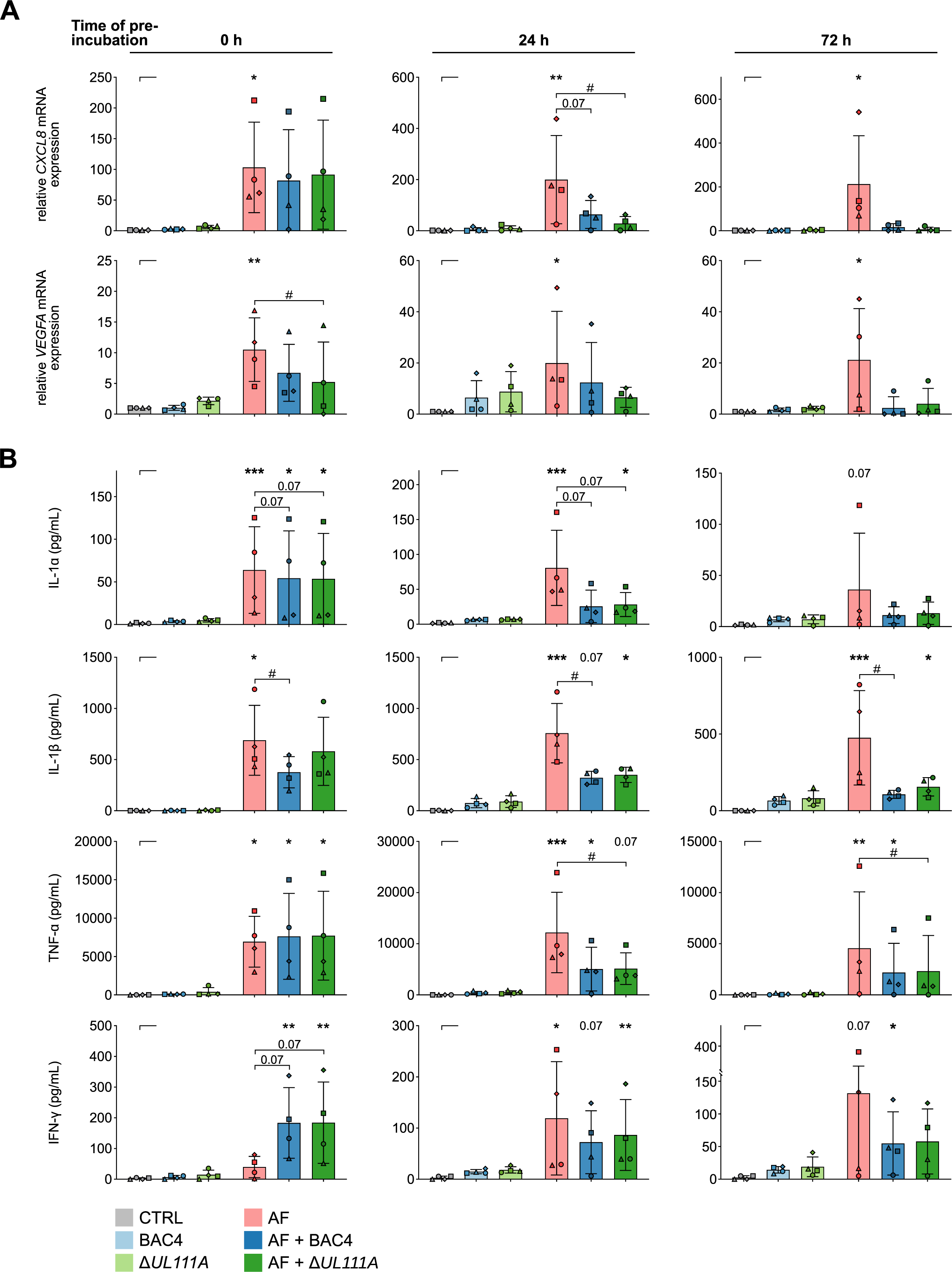
Pre-exposure to HCMV time-dependently weakens antifungal effector responses of moDCs. Relative mRNA expression (A) and cytokine secretion (B) of moDCs (CTRL) (pre)-exposed (0 h, 24 h, 72 h) to BAC4, Δ*UL111A*, and/or *A. fumigatus* (AF), either individually (BAC4, Δ*UL111A*, AF) or in combination (AF + BAC4, AF + Δ*UL111A*). N = 4 independent donors. Columns and error bars indicate means and standard deviations, respectively. Repeated-measures (RM) one-way analysis of variance (ANOVA) with Dunnett’s post-hoc test (A) and Friedman test with Dunn’s multiple comparisons (B) versus “CTRL”, i.e., uninfected moDCs (asterisks). In addition, single AF infection was compared to co-infection (AF + BAC4, AF + Δ*UL111A*) using RM 1-way ANOVA and Dunnett’s post-hoc test (A) and Friedman test with Dunn’s multiple comparisons test (B) versus “AF” (hash signs). */# p < 0.05, **/## p < 0.01, ***/### p < 0.001.

### Interference of HCMV with antifungal responses is largely independent of _CMV_IL-10 expression

We then sought to put these findings into a more global immunobiological context and obtain an unbiased characterization of the HCMV effect on anti-*A. fumigatus* responses specifically in the co-infection setting. To that end, we performed additional analysis with DESeq2 using more complex (single infection corrected) test models for the impact of _CMV_IL-10 competent BAC4, incompetent Δ*UL111A* mutant, or both HCMV strains during co-infection (see Methods). The resulting DEGs were imported into Ingenuity Pathway Analysis to perform three distinct core analyses for HCMV effects in the co-infection setting, considering data for (i) both HCMV strains combined, (ii) WT (BAC4) HCMV, and (iii) the Δ*UL111A* mutant (Figure 6A).

**Figure 6:**
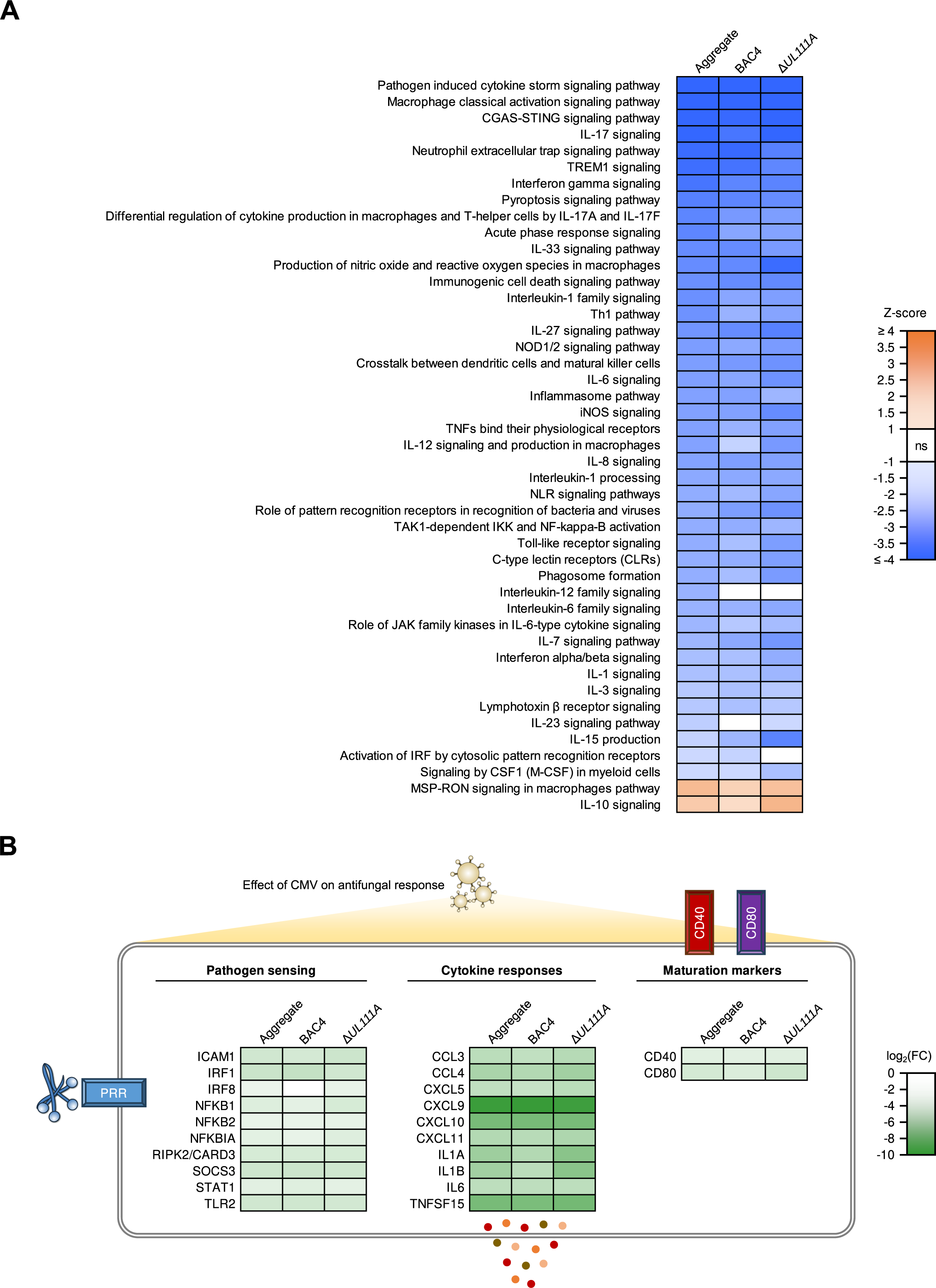
HCMV disrupts antifungal immune responses largely independent of viral _CMV_IL-10 expression. (A) Pathway enrichment analysis to assess the viral single infection-adjusted effect of HCMV on transcriptional responses against *A. fumigatus* during co-infection. Immune-related pathways with absolute z-scores ≥ 1 and Benjamini-Hochberg-adjusted p-values <0.05 are shown. (B) Viral single infection-adjusted expression data for selected individual genes associated with pathogen sensing, cytokine and chemokine production, and moDC maturation to assess the effect of HCMV on antifungal responses during co-infection. (A-B) Three separate analyses were performed, considering data for both HCMV strains on aggregate (left column), WT (BAC4) HCMV only (center column), and the Δ*UL111A* mutant only (right column).

Infection with either HCMV strain significantly suppressed numerous pathways associated with immune recognition of fungal antigens (e.g., Toll-like receptor signaling), inflammasome and acute phase response signaling, key antifungal effector cytokine responses (e.g., IFN-γ, IL-6, IL-8, and IL-17 signaling), activation and maturation of mononuclear phagocytes, and intercellular communication during co-infection (Figure 6A). Inversely, Macrophage Stimulating Protein/*Receptor d’Origine Nantais* (MSP-RON) signaling, a pathway commonly associated with M2 macrophage differentiation, and IL-10 signaling were significantly induced by HCMV in the co-infection setting (Figure 6A). Of note, only 2 out of 43 significantly suppressed immune-related pathways differed significantly between BAC4 and Δ*UL111A* (co-)infection. “Activation of IRF by cytosolic pattern recognition receptors” was the only pathway significantly suppressed by BAC4 (WT) but not by the Δ*UL111A* mutant in the co-infection context (Figure 3A).

On an individual gene level, we found significant single-infection-corrected HCMV-induced suppression of genes associated with pattern recognition receptor and downstream signaling (2.0 – 6.0-fold downregulation on aggregate), cytokine and chemokine transcripts (7.3 – 760.3-fold downregulation), and genes encoding DC maturation markers (2.7 – 4.2-fold downregulation) in the co-infection setting (Figure 6B). Corroborating the pathway-level analysis, _CMV_IL-10 was not essential for significant HCMV-induced downregulation of the mentioned individual transcripts in the co-infection setting (Figure 6B). Collectively, when accounting for expression levels in single HCMV infections, these results suggest a marginal impact of _CMV_IL-10 expression on HCMV’s suppression of anti-*A. fumigatus* responses during co-infection.

### Pre-exposure of moDCs to HCMV weakens moDC-mediated inhibition of fungal growth, independent of _CMV_IL-10

Lastly, we aimed to test whether HCMV or recombinant _CMV_IL-10 directly influence the anti-fungal activity of moDCs. Therefore, we assessed growth and morphogenesis of *A. fumigatus* in the presence of moDCs, with and without simultaneous challenge with BAC4, Δ*UL111A*, or recombinant _CMV_IL-10 (Figure 7A-B). In an alternative scenario, moDCs were pre-exposed to HCMV or _CMV_IL-10 for 24 h or 72 h prior to challenge with *A. fumigatus* (Figure 7A). Hyphal length and branching were quantified using the IncuCyte NeuroTrack feature.

**Figure 7:**
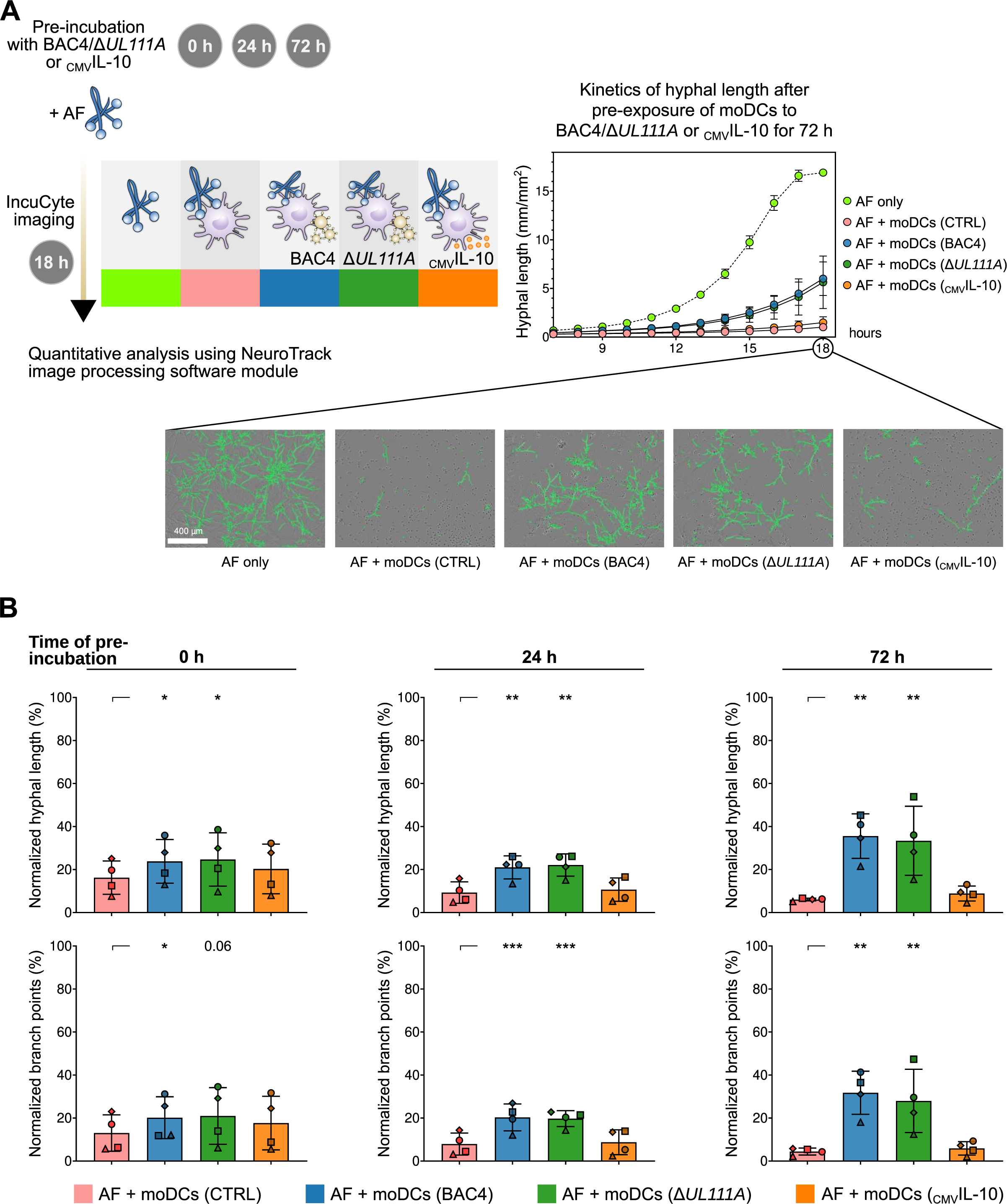
Preincubation of moDCs with HCMV but not with _CMV_IL-10 diminishes the capacity of moDCs to control fungal growth. (A) Left panel: Study design illustrating experimental setup for IncuCyte-based monitoring of fungal growth and branching of co-infection experiments. Right panel: Representative kinetics of *A. fumigatus* (AF) hyphal elongation during transition from germ tubes to hyphae (7–18 h of culture) when cultured alone (AF only) or in the presence of moDCs (AF + moDCs, CTRL), including moDCs pre-exposed for 72 h to BAC4, Δ*UL111A*, or _CMV_IL-10. In addition, images from a representative donor are shown. Green overlays indicate hyphal detection by the NeuroTrack algorithm. (B) Growth and morphogenesis of AF during co-culture with unchallenged moDCs (CTRL) or moDCs pre-exposed for different periods (0 h, 24 h, and 72 h) to moDCs to BAC4, *UL111A*, or _CMV_IL-10. Results are normalized to AF only (= 100%). N = 4 independent donors. Columns and error bars indicate means and standard deviations, respectively. Repeated-measures one-way analysis of variance with Dunnett’s post-hoc test versus “AF + CTRL” (asterisks). * p < 0.05, ** p < 0.01, *** p < 0.001.

Expectedly, naïve moDCs significantly inhibited fungal growth and morphogenesis (Figure 7A-B). Simultaneous challenge of moDCs with _CMV_IL-10 and *A. fumigatus* did not affect moDC-mediated fungal growth inhibition, even after extended pre-exposure (Figure 7B). In contrast, concurrent exposure of moDCs with either HCMV variant modestly reduced their antifungal capacity (mean normalized hyphal length: control 16.2 % vs. BAC4 23.8 % vs. Δ*UL111A* 24.7%; Figure 7B). Additional pre-exposure of moDCs to BAC4 or Δ*UL111A* for 24 h (9.3 % vs. 21.7 % vs. 22.0 %) or 72 h (6.0 % vs. 35.6 % vs. 33.4 %) further weakened the antifungal activity of moDCs, independent of the HCMV variant’s _CMV_IL-10 (Figure 7B). These findings further underscore the limited role of _CMV_IL-10 in anti-*Aspergillus* response and confirm the profound immunosuppressive effect of HCMV on antifungal responses, particularly after prolonged pre-incubation.

## Discussion

HCMV is a highly adaptable pathogen that evades immune surveillance through multiple immunomodulatory mechanisms, with _CMV_IL-10 being considered the preeminent factor in suppressing host immunity (23, 24, 29). Several studies have demonstrated that _CMV_IL-10 downregulates proinflammatory pathways, inhibits antigen presentation, and skews immune responses towards a tolerant state, favoring viral immune escape and potentially exacerbating the risk of secondary infections (24, 26–29). However, the specific role of _CMV_IL-10 in suppressing antifungal immunity has been scarcely studied. To close this knowledge gap, we analyzed immune interferences between HCMV or recombinant _CMV_IL-10 and *A. fumigatus*, comparing the WT virus with a _CMV_IL-10-deficient mutant or _CMV_IL-10.

Although a small cohort study detected _CMV_IL-10 concentrations ranging from 31 to 547 pg/mL in the plasma of (most) HCMV-seropositive healthy donors (37), _CMV_IL-10 levels during lytic infection or reactivation are unknown and are likely significantly higher. Using high-concentration (50 ng/mL) pretreatment of moDCs with _CMV_IL-10 according to a published protocol (27), our data corroborated the protein’s immunosuppressive role, including suppression of the NF-κB–responsive genes *TNF-α* and *IL-1β* (26), interference with *Aspergillus*-induced DC maturation and antigen-presentation (CD40, CD86) (27), and upregulation of the viral entry target CD209 (38) (Figure 8). Moreover, we found enhanced _CMV_IL-10-mediated cellular IL-10 production, which was further amplified in the presence of *A. fumigatus*, contributing to an anti-inflammatory environment (24, 34). However, even high-concentration pretreatment with recombinant _CMV_IL-10 had a minimal impact on fungal clearance in our *in-vitro* assay.

**Figure 8:**
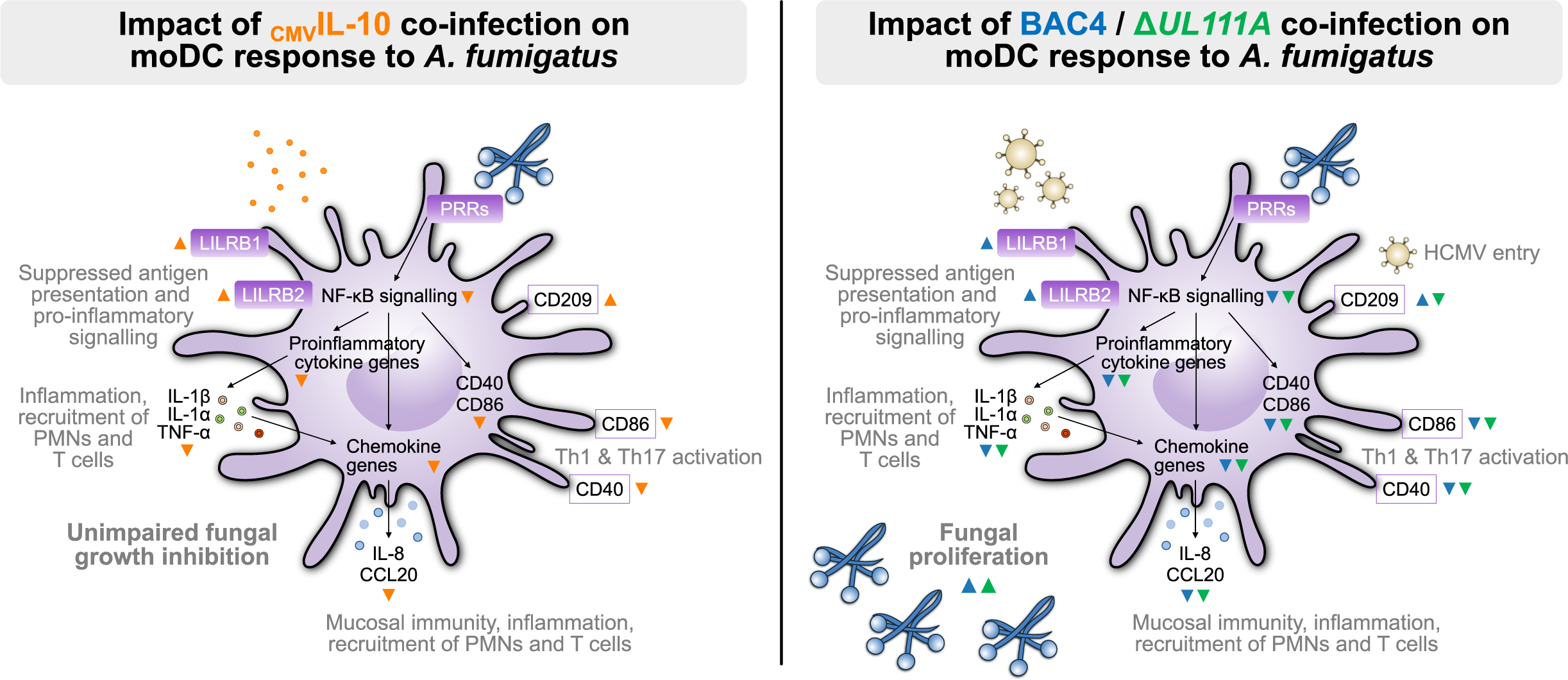
Co-infection with HMCV attenuates various. *A. fumigatus*-induced moDC effector responses in a largely _CMV_IL-10-independent manner. Left part: Orange arrowheads indicate _CMV_IL-10 effects on *A. fumigatus*-induced moDC responses. Right part: Blue and green arrowheads indicate BAC4 and Δ*UL111A* effects on *A. fumigatus*-induced moDC responses. Abbreviations: CCL, CD, Cluster of Differentiation; CCL, C-C motif chemokine ligand; NF-κB, nuclear factor ‘kappa-light-chain-enhancer of activated B cells’; PMN, polymorphonuclear leukocytes; T, T lymphocyte; Th, T-helper cell; LILRB, leukocyte immunoglobulin-like receptor subfamily B; PRR, pattern recognition receptor; IL, interleukin; TNF-α, tumor necrosis factor alpha.

Consistent with our previous findings, HCMV infection upregulated host genes associated with immune clearance, including *CXCL9–11* and genes for type I interferons (IFNA/B) (16, 39–44). While upregulation of cGAS and TLR3 was comparable between BAC4 and Δ*UL111A* infected moDCs, the _CMV_IL-10-deficient Δ*UL111A* mutant induced stronger *IFNA/B* expression and IFN-α/β release, confirming published data (28, 29, 45). Moreover, moDCs infected with Δ*UL111A* showed a trend toward stronger induction of IFN-γ–inducible chemokines and pro-inflammatory cytokines (26, 40, 46), consistent with previous findings that _CMV_IL-10 dampens antiviral responses and promotes an anti-inflammatory environment (24, 29, 34).

Co-infection with either BAC4 or the Δ*UL111A* mutant attenuated various *A. fumigatus*-induced DC effector responses, including pathogen sensing, DC maturation, and cytokine release. Specifically, production of key proinflammatory cytokines (e.g., IL-1α and IL-1β), chemokines (e.g., CCL20 and CXCL8), and VEGFA were markedly downregulated during co-infection with either HCMV variant, reflecting the known synergistic interference between the two pathogens (16). These cytokines and chemokines are essential for recruiting neutrophils and monocytes, reinforcing epithelial barriers, supporting tissue remodeling, and promoting type-17 T-helper cell responses (16, 47–53). Disruption of these responses might collectively result in profoundly compromised antifungal immunity during co-infection.

Interestingly, both *LILRB1* and *LILRB2* were markedly upregulated in response to HCMV infection, especially after BAC4 infection, with elevated expression already evident in BAC4 single infection and further amplified during co-infection. Accordingly, moDCs pre-exposed to recombinant _CMV_IL-10 and subsequently infected with *A. fumigatus* showed enhanced transcription of *LILRB1* and *LILRB2*. These inhibitory receptors are known to attenuate antigen-presenting cell activation and suppress proinflammatory signaling (35, 36), pointing to a _CMV_IL-10–dependent mechanism by which BAC4 may modulate host immune responses. However, our observation of overall high functional similarity between Δ*UL111A* and BAC4 in suppressing antifungal immunity aligns with the modest impact of _CMV_IL-10 pre-exposure on the anti-*Aspergillus* response, suggesting an overall minor impact of _CMV_IL-10 expression on antifungal immunity in the co-infection context.

A strength of our study has been the assessment of immune dynamics after different pre-exposure periods of moDCs to HCMV. The virus follows a tightly regulated replication cycle with sequential phases of gene expression (54–56). Of note, proteins involved in host immune interference are expressed rapidly in the viral replication cycle (within the first few hours) (54, 55, 57, 58). However, many structural proteins required for virion assembly are produced in the later stages (> 24 hours), resulting in the release of a huge number of infectious virions between 48 and 72 h (57, 59, 60). Thus, pre-incubation of moDCs with either HCMV variant for 24 h or 72 h and subsequent *A. fumigatus* challenge led to even stronger suppression of key cytokine responses and the DC’s ability to control fungal growth, regardless of _CMV_IL-10 expression (34, 61).

Importantly, interferences with antifungal immunity are not exclusive to HCMV but have been documented extensively in patients and preclinical models with various viral infections, especially influenza or COVID-19 (62–66). Thus far, the relative impact of host immune dysregulation versus the co-pathogens’ specific virulence factors on antifungal immune failure during co-infections remains poorly understood. Although different underlying viral infections were associated with considerable differences in the acuity and severity of antifungal immune paralysis and its specific cellular and humoral determinants, there is increasing evidence for commonalities in viral dysregulation of host responses (19, 64, 66). Two commonly reported denominators of antifungal immune compromise during viral co-infections are the impairment of pathogen sensing pathways and dysregulated cytokine signaling, including both cytokine-mediated immunotoxicity (e.g., due to type-1 and 2 interferons) and severe attenuation of key antifungal effector cytokines (15, 16, 62, 67). These common denominators of virus-induced immune paralysis were also seen during HCMV co-infection in the present study, regardless of _CMV_IL-10 expression. Given the significant immunologic commonalities across various viral and fungal co-infections despite the co-pathogens’ vastly dissimilar biology, the limited contribution of a singular viral virulence factor exclusively present in HCMV seems conceptually plausible.

However, _CMV_IL-10 might function within a broader network of viral immune evasion proteins. Beyond _CMV_IL-10, HCMV encodes several other immunosuppressive proteins that may contribute to an increased risk of fungal superinfections (3, 68). Specifically, US2, US3, US6, and US11 can interfere with antigen presentation by downregulating MHC class I and II molecules (69–72). Additionally, US28 modulates DC migration through altered chemokine signaling (73, 74). Lastly, UL23 was shown to attenuate IFN-γ signaling and exploit the PD-L1 immune checkpoint pathway to attenuate adaptive host immunity (75). Possibly synergizing with _CMV_IL-10, these immune evasion strategies might cause pleiotropic breaches to antifungal immunity. Thus, the role of these immunosuppressive HCMV proteins in antifungal immunity should be analyzed in future studies using defined CMV mutants and additional immune cell populations such as T cells (27).

Furthermore, our moDC-focused study did not account for a possible broader impact of _CMV_IL-10 on other immune cell subsets (24). For instance, _CMV_IL-10 modulates NK-cell function by altering cytokine signaling directly or indirectly, influencing their cytotoxic potential against infected cells (24, 27, 76). However, the rapid NK cell-mediated lysis of virally infected cells *in-vitro* would limit the ability to fully recapitulate the complexity of intercellular interactions (77–79). In contrast, moDCs are a more robust and tractable model system that facilitates controlled and reproducible functional analyses (80, 81). Likewise, the role of epithelial cells, which serve as key sites for HCMV entry, replication, and dissemination, was not accounted for in our study (82). Epithelial cells not only support viral persistence but can also contribute to local immune dysregulation through cytokine release and direct crosstalk with infiltrating immune cells (82–85). Thus, future studies should characterize the impact of HCMV and _CMV_IL-10 on epithelial barrier integrity, cytokine signaling, and crosstalk between innate and adaptive immune compartments, as they relate to antifungal defense.

Relying on cells from healthy donors, our study also did not capture critical co-variables (e.g., transplant history, immunosuppressive pharmacotherapy, other underlying conditions, and co-infections) that could significantly influence viral pathogenesis and immune modulation in immunocompromised hosts (8, 86, 87). However, the large blood volumes required for generation of a sufficient number of moDCs have precluded assessment of patient cells in the present study (88). Alternatively, *in-vivo* models incorporating both underlying immunosuppression (e.g., agents used for prophylaxis or therapy of graft-versus-host disease) and HCMV infection could provide further mechanistic insights into the role of _CMV_IL-10 in antifungal immunity. However, the strict host specificity of HCMV would require the use of advanced humanized mouse models that are intricate, expensive, and insufficiently validated for studies of invasive aspergillosis (89–91).

In summary, our data underscore the rapid and pleiotropic impact of HCMV on DCs, a key cellular player in anti-*A. fumigatus* immunity. _CMV_IL-10 exerted a strong immunosuppressive effect on moDCs by downregulating proinflammatory pathways, especially IFN-γ-inducible chemokine production, type I interferon secretion, and antigen presentation. However, despite its role as a master orchestrator of host immune evasion, the impact of _CMV_IL-10 on antifungal defense during co-infection was limited. Overall, this points to a complex interplay of immunological, genetic, environmental, and iatrogenic factors that collectively drive co-infection risk in HCMV-infected individuals. Future research should delineate the broader immunoregulatory effects of HCMV to inform the development of therapeutic strategies to restore immune homeostasis in high-risk patient populations and reduce the burden of co-infections in immunocompromised individuals with HCMV infection.

## Methods

### Primary cell isolation

MoDCs were generated from CD14^+^ monocytes isolated from peripheral blood mononuclear cell as previously described (16, 80). Processing of human peripheral venous blood from healthy adults was approved by the Ethics Committee of the University Hospital Würzburg (#302/12).

### Growth condition of *A. fumigatus* and preparation of germlings

*A. fumigatus* strain American Type Culture Collection (ATCC) 46645 or *Aspergillus* reporter (FLARE) conidia (92) were cultured on beer wort agar at 35 °C until conidiophore formation. Conidial suspensions were harvested using sterile water and filtered through a 20-µm cell strainer. To induce germ tube formation, 2×10^7^ conidia (ATCC46645) were incubated in RPMI medium while shaking (200 rpm, 25 °C) until small germlings became visible. Germlings were centrifuged at 5,000 ×*g* for 10 min and resuspended in CellGenix medium (2×10^7^/mL).

### Generation of Δ*UL111A*

TB40-BAC-KL7-SE-EGFP-*UL111A*STOP was generated using the markerless mutagenesis protocol (93). In brief, a recombination fragment was generated by PCR from plasmid pEP-Kan-S with primers: 5′-GGAGGCGAAGCCGGCGACGACGACGACGATAAAGAATACATGACCGCAGTGTCGTTAGGAGGATTACGCGACCAGATTAGGATGACGACGATAAGT-3′ and 5′-AGGTGACGCGGAGATCTTGCAATCTGGTCGCGTAATCCTCCTAACGACACTGCGGTCATGTATTCTTTAT CGTCGTCGCAACCAATTAACCAATTCTGA-3′ and electroporated into recombination-activated GS1783 bacteria harboring TB40-BAC-KL7-SE-EGFP (94). After removal of the selection marker, BAC DNA was isolated using the NucleoBond Xtra Midi Kit (Macherey-Nagel) and the integrity of the DNA was checked by restriction fragment length analysis (RFLA) and sequencing. BAC DNA was transfected into human foreskin fibroblasts with the K2 Transfection System (Biontex). When cytopathic effect reached 100%, supernatants were harvested by centrifugation at 3200 ×*g* for 10 min to remove cellular debris and subsequently frozen at -80 °C.

### moDC infection assay

Primary human moDCs were infected with HCMV strains TB40-BAC-KL7-SE-EGFP (94) or TB40-BAC-KL7-SE-EGFP-*UL111A*STOP (Δ*UL111A*, lacking _CMV_IL-10) at a multiplicity of infection (MOI) of 3 and centrifuged at 300 ×*g* for 30 min to initiate proper HCMV infection. Alternatively, moDCs were exposed to recombinant _CMV_IL-10 (50 ng/mL, R&D Systems) for 24 h. For subsequent fungal infection, *A. fumigatus* germlings were added at an MOI of 0.5.

For flow cytometric analysis, moDCs were pre-incubated with HCMV or _CMV_IL-10 for 24 h. Subsequently, the cells were incubated with *A. fumigatus* germlings for an additional 9 h.

### RNA isolation and bulk transcriptomic profiling

MoDCs cells were harvested and centrifuged (300 ×*g*, 10 min). Cell pellets were frozen in RNAprotect Cell Reagent (Qiagen) at -80°C until further analysis. RNA integrity was determined with a 2100 BioAnalyzer (Agilent Technologies). RIN values of all samples were above 8.5.

Library preparation was performed with the Illumina TruSeq Stranded mRNA technology, according to the manufacturer’s protocol. RNA sequencing was performed by IMGM Laboratories GmbH (Martinsried) on the Illumina NovaSeq® 6000 with 1×100-150 bp single-read chemistry. Raw files are accessible at ArrayExpress (https://www.ebi.ac.uk/biostudies/arrayexpress), accession number E-MTAB-15110.

### RNA sequencing data processing

Preprocessing of raw reads, including quality control and gene abundance estimation, was done with the GEO2RNaseq pipeline (v0.9.12, (95)) in R (version 3.5.1). Quality analysis was done with FastQC (v0.11.8) before and after trimming. Read-quality trimming was done with Trimmomatic (v0.36). Reads were rRNA-filtered using SortMeRNA (v2.1) with a single rRNA database combining all rRNA databases shipped with SortMeRNA. Reference annotation was created by extracting and combining exon features from corresponding annotation files. Reads were mapped against the reference genome of *A. fumigatus* (Af293, ASM265v1) using HiSat2 (v2.1.0, single-end mode). Gene abundance estimation was done with featureCounts (v1.28.0) in single-end mode with default parameters. MultiQC version 1.7 was finally used to summarize and assess the quality of the output of FastQC, Trimmomatic, HiSat, featureCounts, and SAMtools. The count matrix with gene abundance data with and without median-of-ratios normalization were extracted (96). Statistical analysis and applied analytic models are summarized in Supplementary Methods.

### Ingenuity Pathway Analysis

Normalized expression levels of genes with a significant single-infection-corrected HCMV effect in the co-infection setting (p-value < 0.05 and false discovery rate < 0.2) were imported into Ingenuity Pathway Analysis (Qiagen). Core analysis was performed to determine canonical pathway enrichment. Pathway enrichment was considered significantly different at an absolute z-score value ≥ 1 and Benjamini-Hochberg adjusted p value < 0.05. Individual genes associated with pathogen sensing, cytokine responses, and dendritic cell maturation were selected from the top 20 immune-related pathways suppressed by HCMV co-infection (lowest z-scores).

### cDNA synthesis and quantitative reverse transcription PCR

To validate differential gene expression, total RNA, extracted with the RNeasy Plus Mini Kit (Qiagen) according to the manufacturer’s instructions, was converted into first-strand cDNA using the First Strand cDNA Synthesis Kit (Thermo Fisher Scientific). qRT-PCR reactions were set up in duplicates using 10 mM gene-specific primers and SYBR GreenER™ qPCR SuperMix Universal (BioRad Laboratories) on a StepOnePlus™ Real-Time PCR System (Applied Biosystems). Relative gene expression was calculated using the ΔΔCt method relative to a house-keeping gene (GAPDH). Primers are summarized in Table S1.

### Flow cytometry

Cells were harvested, washed (5 min at 500 ×*g*), and resuspended in the antibody solution for extracellular staining along with the fixable Viobility^TM^ Live/Dead Dye (Miltenyi Biotec). Intracellular staining was performed using the Inside Stain Kit (Miltenyi Biotec) according to the manufacturer’s instructions. Antibodies, clones, and manufacturers are summarized in Table S2. Flow cytometric data was acquired on a Cytoflex (Beckman Coulter) and analyzed using Kaluza v2.1 software (Beckman Coulter). The gating strategy is summarized in Figure S3.

### Multiplex assays for cytokine secretion

For multiplexed quantification of cytokine and chemokine concentrations (IFN-α, IFN-γ, IL-1α, IL-1β, IL-2, IL-6, IL-8, IL-10, IL-23, CXCL10, CXCL11, CCL2, CXCL9, CCL3, CCL4, CCL20, CCL5, and TNF-α (cells used for RNA-Seq) or IFN-α, IFN-β, IFN-γ, IL-1α, IL-1β, IL-2, IL-6, IL-8, IL-10, IL-12p70, IL-17A, IL-23, CXCL10, CXCL11, CCL5 and TNF-α (cells time-dependently exposed to HCMV) in supernatants of stimulated moDCs, a ProcartaPlex 18-PLEX assay kit (ThermoFisher Scientific) was used according to the manufacturer’s protocol. Data acquisition and analysis were performed using a Bio-Plex 200 Luminex reader in combination with Bio-Plex Manager Software version 6.2 (Bio-Rad).

### IncuCyte time-lapse imaging

To test the immunomodulatory capacity of HCMV on the direct antifungal response of moDCs, *A. fumigatus* FLARE conidia were suspended in CellGenix medium (4×10^3^ conidia/mL). Fifty microliters (200 conidia/well) were dispensed per well of a 96-well flat-bottom plate. Ten thousand moDCs (effector/target ratio, 50) that had been pre-incubated with BAC4 or Δ*UL111A* (MOI 3), _CMV_IL-10 (50 ng/mL), or plain medium for 0, 24, or 72 h were added in 100 μL CellGenix medium. IncuCyte microscopy and NeuroTrack-based image analysis were performed following a published protocol (97), as detailed in Supplementary Methods.

### Statistical analyses

Microsoft Excel 365, GraphPad Prism version 10, and R version 3.5.1 were used for data compilation, analysis, and visualization. Depending on the data format, significance testing was performed using Friedman test with Dunn’s multiple comparisons test, paired Wilcoxon signed-rank test, repeated measures one-way analysis of variance (ANOVA) with Dunnett’s post-test, or paired t-test. Significance tests are specified in the figure legends. (Adjusted) p-values < 0.05 were considered significant.

## Author Contributions

The study was conceived by LH, LB, SW, SaS and JL. Experiments were planed and performed by LH, LB, LS, KH, OK, SW and JL. Resources were provided by AG, and LD, KLS. Data were analyzed by LH, LB, SS and SW. Data were visualized by LH, LB, SS and SW. Project administration and supervision were led by OK, LD, GP, AJW, HE, SW, SaS and JL. Funding was acquired by GP, HE and JL. The original draft was written by LH, LB, SW, SaS and JL. All co-authors reviewed, edited, and approved the manuscript.

## ACKNOWLEDGMENTS

The study was supported by the Deutsche Forschungsgemeinschaft (DFG) within the Collaborative Research Center CRC TR124 FungiNet “Pathogenic fungi and their human host: Networks of interaction”, DFG project number 210879364 (project A2 to HE and JL, C3 to OKu, and INF to GP) and within the CRC SFB 1586 DECIDE „DECisions in Infectious DisEases“(#492620490; project A04 to AJW, A06 to JL, A07 to LD, and C03 to OKu). We thank Prof. Tobias M. Hohl, (Infectious Disease Service, Department of Medicine, Memorial Sloan Kettering Cancer Center, New York, NY, USA) and Prof. Andreas Beilhack (Department of Internal Medicine II, University Hospital Würzburg, Center of Experimental Molecular Medicine, Würzburg, Germany) for providing *A. fumigatus* FLARE conidia. The authors also thank all blood donors.

