## Supplementary Figures S1-S3, Supplementary Tables for "Unveiling immune interference: How the dendritic cell response to co-infection with *Aspergillus fumigatus* is modulated by human cytomegalovirus and its virokine CMVIL-10"

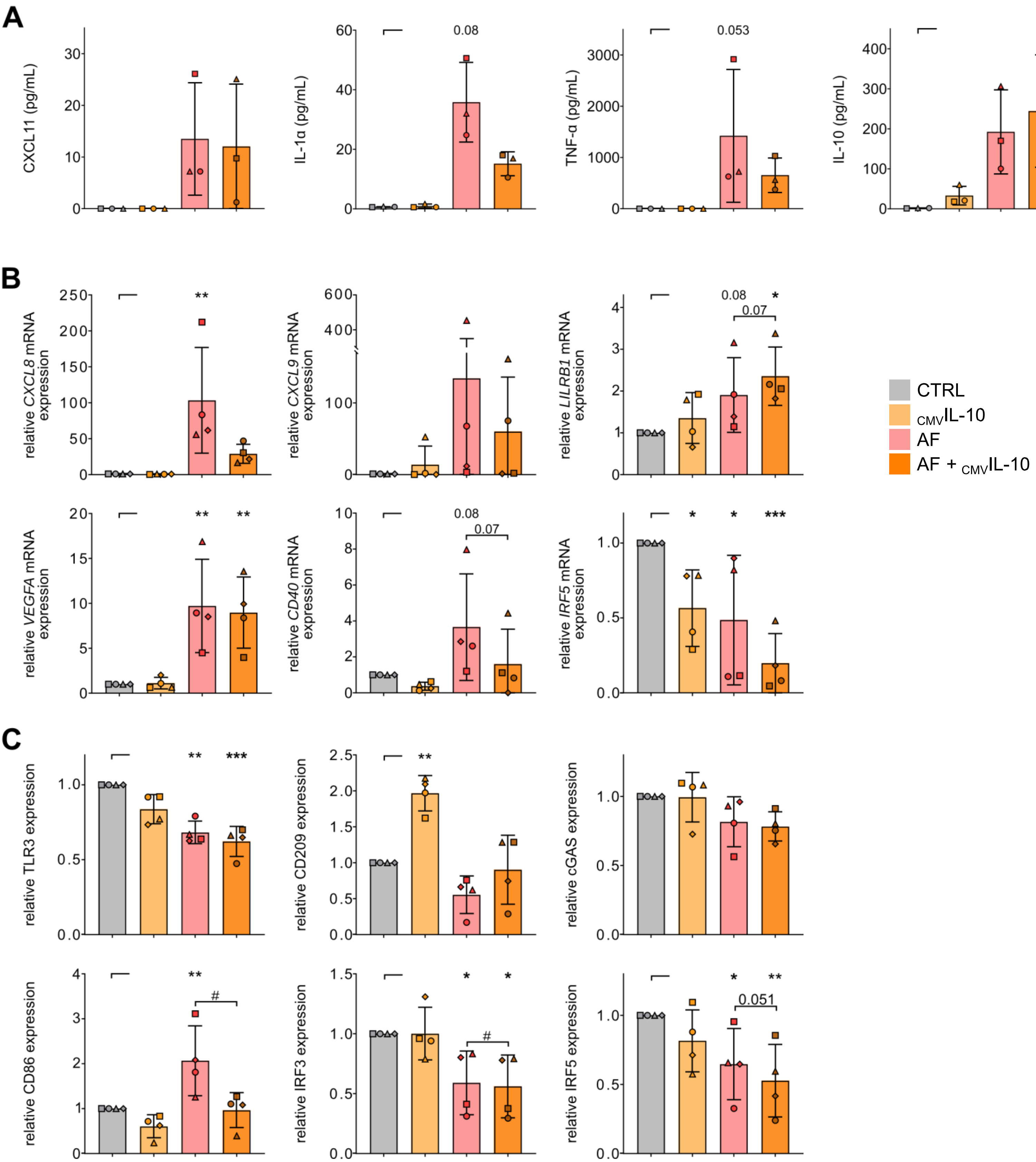

**Figure S1: Recombinant $_{CMV}$IL-10 does not significantly alter *A. fumigatus*-induced responses of moDCs at both mRNA and protein levels.**

Cytokine/chemokine release (A), relative mRNA expression (B), and flow cytometric analysis (C) of naïve moDCs (CTRL) pre-incubated with $_{CMV}$IL-10 for 24 h prior to 9 h culture with or without *A. fumigatus* (AF) infection. (A) N = 3 independent donors. (B-C) N = 4 independent donors. (A-C) Columns and error bars indicate means and standard deviations, respectively. (A) Friedman test with Dunn's multiple comparisons test versus CTRL (asterisks). (B-C) Repeated measures one-way analysis of variance with Dunnett's post-hoc test versus "CTRL", i.e., uninfected moDCs (asterisks). In addition, single AF infection was compared to co-stimulation (AF + $_{CMV}$IL-10) using paired t-Test (hash signs). */# $p < 0.05$, **/## $p < 0.01$, ***/### $p < 0.001$.

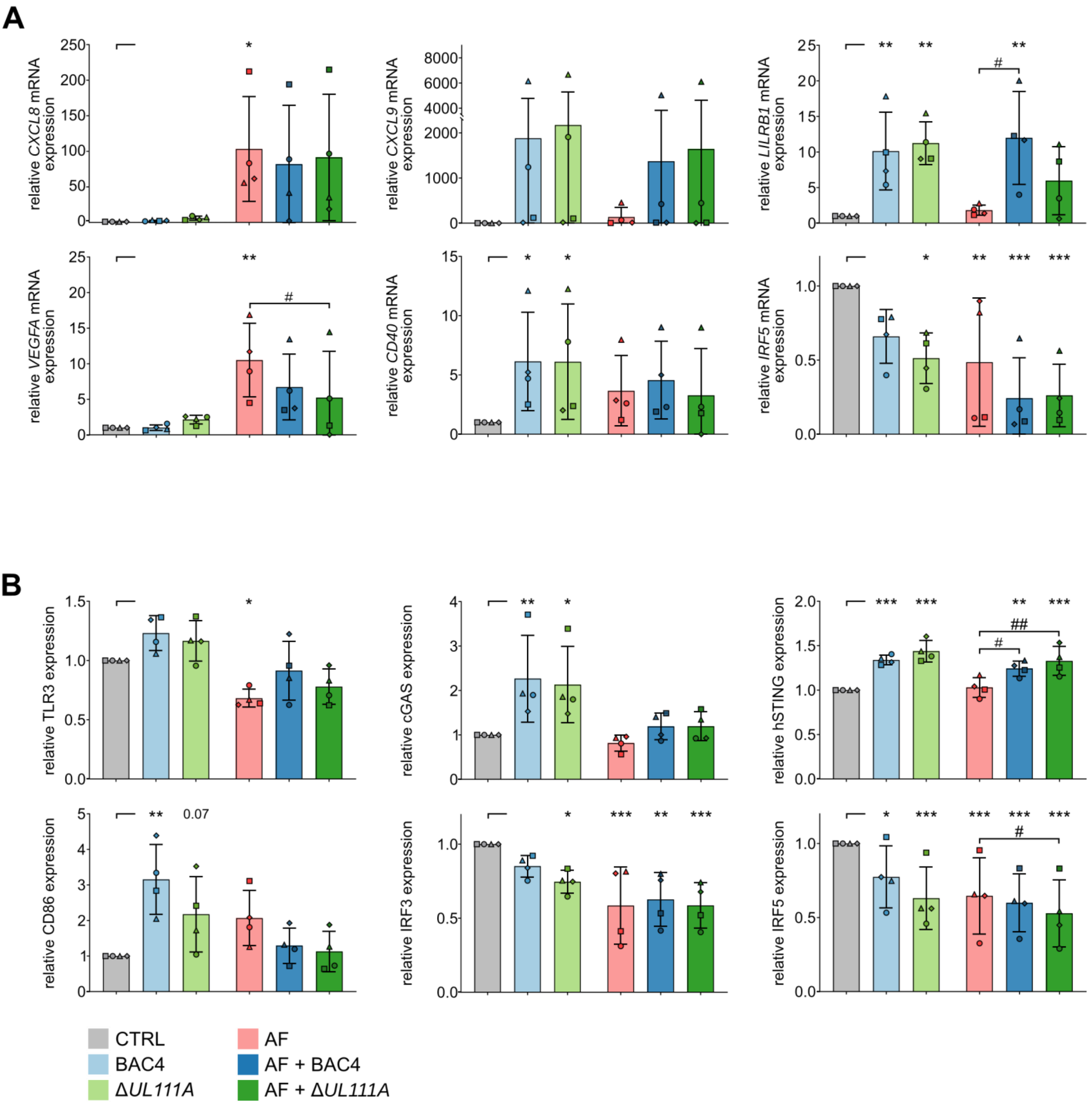


**Figure S2: $_{CMV}$IL-10 modulates the antiviral response of moDCs but has minimal effects on HCMV-mediated suppression of antifungal responses at both mRNA and protein levels.**

(A) Relative mRNA expression of moDCs confronted with BAC4, Δ*UL111A*, and/or *A. fumigatus* (AF), either individually (BAC4, Δ*UL111A*, AF) or in combination (AF + BAC4, AF + Δ*UL111A*). (B) Flow cytometric analysis of moDCs pre-incubated with BAC4 or Δ*UL111A* for 24 h prior to culture with or without AF infection for 9 h. (A-B) N = 4 independent donors. Columns and error bars indicate means and standard deviations, respectively. Repeated measures (RM) one-way analysis of variance (ANOVA) and Dunnett's post-hoc test versus "CTRL", i.e., uninfected moDCs (asterisks). In addition, single AF infection was compared to co-infection (AF + BAC4, AF + Δ*UL111A*) using RM one-way ANOVA and Dunnett's post-hoc test versus "AF" (hash signs). */# $p < 0.05$, **/## $p < 0.01$, ***/### $p < 0.001$.

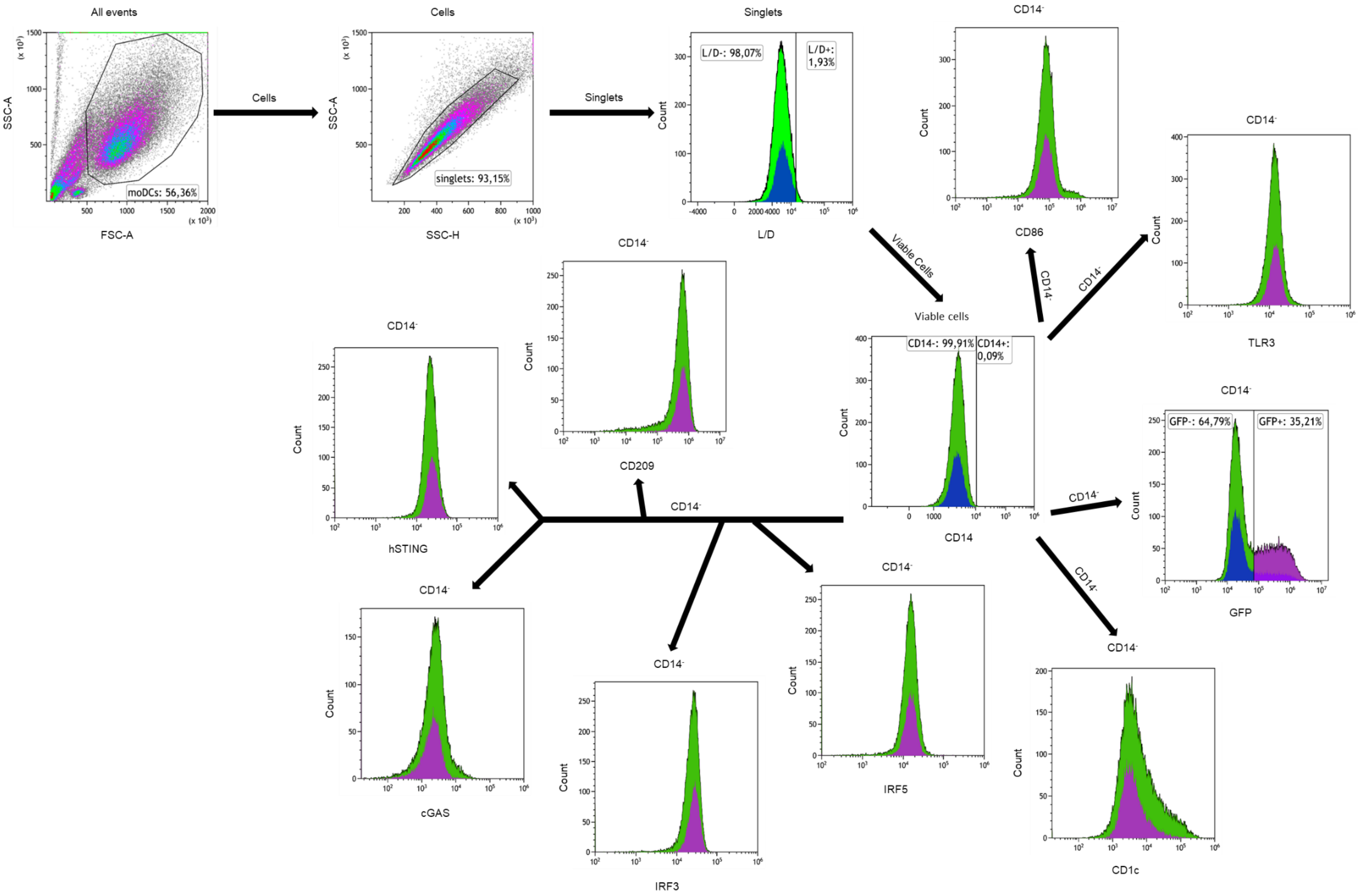

All events
moDCs: 56,36%
SSC-A
FSC-A
Cells
singlets: 93,15%
SSC-H
Singlets
L/D-: 98,07%
L/D+: 1,93%
Count
L/D
Viable Cells
Viable cells
CD14-: 99,91%
CD14+: 0,09%
CD14
CD14-
CD86
TLR3
GFP-: 64,79%
GFP+: 35,21%
GFP
CD1c
IRF5
IRF3
cGAS
hSTING
CD209

**Figure S3. Gating strategy to study efficancy of HCMV infection of moDCs, moDC maturation and activation.**

MoDCs were selected according to their forward and side scattering and single myeloid cells were gated based on SSC-A, and SSC-H properties. Dead moDCs were excluded by Live/Dead staining. moDCs were identified as $CD14^-$. $GFP^+$ cells were defined as HCMV infected moDCs. Mean fluorescence intensities (MFI) for moDC cell surface markers and intracellular markers showing modulations after HCMV or *A. fumigatus* challenge (CD1c, CD86, TLR3, IRF3, IRF5, cGAS, hSTING, CD209) were gated within each $CD14^-$ moDC cell population. MFI was determined for all markers and calculated in comparison to the MFI of their corresponding unstimulated moDCs control (CTRL).

**Supplementary Tables:**

Table S1: Primers used for quantitative real-time PCR

| | | |
|---|---|---|
| **CXCL8** | fw | GTTTTTGAAGAGGGCTGAG |
| | rev | TTTGCTTGAAGTTTCACTGG |
| **VEGFA** | fw | AATGTGAATGCAGACCAAAG |
| | rev | GACTTATACCGGGATTTCTTG |
| **IRF5** | fw | CTCAGCCCTACAAGATCTAC |
| | rev | CTGCACCAAAAGAGTAATCC |
| **CD40** | fw | CTGGCACTGTACGAGTGAGG |
| | rev | AAGACCAGCACCAAGAGGATG |
| **CXCL9** | fw | AGGTCAGCCAAAAGAAAAAG |
| | rev | TGAAGTGGTCTCTTATGTAGTC |
| **LILRB1** | fw | GAATGAGGAGAAAGCAAGAAG |
| | rev | TGAGCTTGATGTAAATGTGC |
| **GAPDH** | fw | GGACTGAGGCTCCCACCTTT |
| | rev | GCATGGACTGTGGTCTGCAA |

Table S2: Flow cytometry antibodies and reagents

| Antibodies | Source | Identifier |
|---|---|---|
| Anti-human CD1c (L161) BrilliantViolet650 | Biolegend, San Diego, CA, USA | Cat#331542 |
| Anti-human CD14 (REA599) Viogreen | Miltenyi Biotec, Bergisch Gladbach, Germany | Cat#130-110-583 |
| Anti-human CD86 (REA968) APC-Vio770 | Miltenyi Biotec, Bergisch Gladbach, Germany | Cat#130-116-163 |
| Anti-human TLR3 (TLR 3.7) APC * | Miltenyi Biotec, Bergisch Gladbach, Germany | Cat#130-096-885 |

| | | |
|---|---|---|
| Anti-human CD209 (REA617) PE-Vio770 * | Miltenyi Biotec, Bergisch Gladbach, Germany | Cat#130-128-219 |
| Anti-human cGAS (E5V3W) Alexa Fluor 647 * | Cell Signaling, Beverly, MA, USA | Cat#43398S |
| Anti-human hSTING (723505) PE * | R&D Systems, Minneapolis, MN, USA | Cat#IC7169P |
| Anti-human IRF3 (482205) Alexa Fluor 750 * | R&D Systems, Minneapolis, MN, USA | Cat#FAB4019S |
| Anti-human IRF5 Alexa Fluor 700 * | R&D Systems, Minneapolis, MN, USA | Cat#IC4508N |

**Supplementary Methods:**

RNA sequencing statistical analysis and analytic models

Differential gene expression analysis was done using DESeq2 (v1.46.0, R version 4.4.2) considering a multiple-test adjusted $p \leq 0.05$ and log2(fold-change) $\geq 1$ as cut-offs for significance. Given donor heterogeneity, DESeq2 statistical designs were created to control for donor origin (design = ~ donor + condition) and different pairwise condition differences, unless otherwise noted. Additional DESeq2 designs to control for single infections when testing co-infection (CoI) with $_{\text{CMV}}$IL-10 competent BAC4 HCMV strains (design = ~ donor + BAC4 + AF + CoI) $_{\text{CMV}}$IL-10 incompetent Δ*UL111A* strain (design = ~ donor + Δ*UL111A* + AF + CoI) or both (design = ~ donor + BAC4 + Δ*UL111A* + AF + CoI) were applied, where all controlling factors except donor were one-hot encoded (control sample = yes or no, BAC4 sample = yes/no, etc.).

IncuCyte time-lapse imaging

In brief, well plates were imaged hourly in an IncuCyte Zoom HD/2CLR time-lapse microscopy system (Sartorius, Göttingen, Germany) equipped with an IncuCyte Zoom 10× Plan Fluor objective (Sartorius, Göttingen, Germany) for a period of 18 h. Acquisition time for the red channel was 300 ms. The following parameters were used for NeuroTrack analysis: neurite coarse sensitivity, 10; neurite fine sensitivity, 0.75; neurite width, 4 µm. Neurite length [mm/mm$^2$] and numbers of branch points [1/mm$^2$]) were compared to an "*A. fumigatus* only" control without moDCs and/or HCMV/ $_{CMV}$IL-10.
